# Microbiota-dependent inflammation promotes metabolic disorder via NF-κB-mediated inhibition of SREBP in *Drosophila* adipocytes

**DOI:** 10.1101/2021.04.28.441756

**Authors:** Bernard Charroux, Julien Royet

## Abstract

Bacteria that colonize eukaryotic gut have profound influences on the physiology of their host. In *Drosophila*, many of these effects are mediated by the adipocytes that combine immune and metabolic functions. We show here that gut colonization by specific bacteria species stimulate lipogenesis in surrounding enterocytes but also in remote fat body cells and ovaries. This bacteria-dependent lipid production is mediated by SREBP and requires a functional insulin signaling. However, it is antagonized by microbiota-born peptidoglycan which by activating NF-κB signaling, cell-autonomously represses SREPP activation in adipocytes but not in enterocytes. We finally show that by reducing microbiota-derived PGN, the gut-produced PGRP-LB amidase balances host immune and metabolic responses of the fat body to gut-associated bacteria. In the absence of such modulation, uncontrolled immune pathway activation prevents lipid production by the fat body resulting in infection-dependent host death.

**Bullets:**

- Gut microbiota activates lipogenesis locally in enterocytes and remotely in adipocytes
- Bacteria-dependent activation of SREBP in adipocytes is mediated by insulin signaling
- Activation of lipid synthesis by microbiota is antagonized by NF-κB signaling triggered by gut-born peptidoglycan
- By reducing peptidoglycan circulating levels, PGRP-LB maintains a balance between immune and metabolic response to microbiota

## Introduction

In order to develop to adulthood and to later survive in their environment, multi-cellular organisms constantly adapt their metabolism needs to the nutrient availability. These nutrients come from food sources that are unavoidably contaminated by microbes on which they proliferate. Some of the microorganisms ingested with food, or those already associated with the digestive tract, directly participate to the host nutrition either by serving as food themselves or by metabolizing ingested aliments. These transient or permanent gut-associated microbes need to be either tolerated by the host if beneficial, or eliminated if detrimental, a function dedicated to the immune system. Hence, metabolism and immunity, that regulate the host’s responses to these environmental inputs, nutriments and microbes, have co-evolved to provide a coordinated output at the organismal level. In mammals, this optimized response benefits from the fact that some immune cells are embedded into the adipose tissue ^1–3^. Immune cells act as direct regulators of fat metabolism and innate immune signaling can impact metabolic responses cell-autonomously or via systemic inflammation ^4–10^. Beside its role in lipid storage and energy expenditure, the adipose tissue is thus considered as an immune organ able to simultaneously sense nutrient and detect microorganism-derived compounds. Communication between the immune cells and adipocytes is essential to coordinate an *ad hoc* host metabolic response in physiological conditions and in response to microbial challenges ^2^.

In *Drosophila*, the fat body is the major site for lipid depository and combines energy storage, de novo synthesis, and breakdown functions that, in vertebrates, are dedicated to adipose and hepatic tissues ^11,12^. In addition, via the production of many immune effectors including antimicrobial peptides, it plays a key role in orchestrating the innate immune responses to microbial infection ^13–15^. Hence, *Drosophila* provides unique advantages to unravel the complex integration and regulation of these two essential physiological systems, before they evolved into more complex organs in vertebrates. Previous work has shown that *Drosophila* infection with bacteria or with the intracellular parasite *Tubulinosema ratisbonensis* leads to a depletion of fat body lipid stores ^16^. Other studies, based on gain-of-function approaches, revealed that ectopic activation of the NF-κB pathways either Toll or IMD can result in lipid storage reduction. More precisely, immune signaling activation shifts anabolic lipid metabolism from triglyceride storage to phospholipid synthesis to support immune function ^17^.

Former results have shown that immune activation in the fat body cells can be triggered by bacteria present in the digestive tract. For that, the bacterial cell wall component peptidoglycan produced by gut-associated bacteria must cross the gut epithelium and reach the circulating hemolymph where it gets in contact with remote tissues. By activating receptors of the PGRP family expressed in adipocytes this gut-born bacterial ligand activates an NF-κB dependent AMP production^18–20^. This effect is buffered by the PGRP-LB amidase that, by cleaving the PGN into non-immunogenic fragments, prevents a diffusion of PGN to the hemolymph and hence a constant deleterious NF-κB activation in fat body cells of orally infected flies ^20,21^.

In the present study, we analyze the coordinate metabolic and immune responses of *Drosophila* to the presence of immunogenic bacteria in the intestine. We show that flies fed with specific bacteria species including *Escherichia coli* (*E. coli*) and *Erwinia carotovora carotovora* (*E.cc*) trigger a lipogenic program both locally in enterocytes and remotely in adipocytes. We present genetic evidence that this effect is mediated by the activation of SREBP and requires a functional insulin signaling pathway. We also show that by activating the NF-κB/IMD pathway in adipocytes, PGN released by the same bacteria, is cell-autonomously antagonizing SREBP-dependent lipogenesis in adipocytes. Finally, we demonstrate that by regulating the levels of circulating PGN via the PGRP-LB amidase, flies can adjust their metabolic and immune responses towards gut bacteria.

## Results

### Specific gut bacteria species activate an SREBP-dependent lipogenesis in adult adipocytes

Our previous data showed that *Drosophila* enteric infection by the phytopathogen *E. cc* affects lipogenesis in adult adipocytes ^20^. To further characterize this phenomenon, we monitored the activation of SREBP, a master regulator of lipogenesis, in response to bacterial gut colonization. We took advantages of the *Gal4::SREBP* reporter whose transcription relies on the native SREBP promoter and in which the resulting chimeric protein is processed like the endogenous SREBP ^22^. We generated a novel *LexA::SREBP* reporter which mimics *Gal4::SREBP* activation (Supplementary Fig.1, 2a, b). Both chimeric fusion proteins are proteolytic cleaved and respectively activate UAS and LexAop fluorescent reporters, in cells wherein SREBP ensures *de novo* lipid synthesis ^22–25^ (Supplementary Fig.1, 2a, b).

In addition to its constitutive activation in oenocytes already reported ^22^, *Gal4::SREBP* expression was detected in fat bodies of flies orally fed with *E.cc* (Fig. 1a, b). When *E. coli* was used to orally infect flies, an even stronger fat body SREBP activation was observed (Fig. 1a, b). Other bacteria species such *Lactobacillus plantarum^WJL^* (*L. plantarum^WJL^*), *Acetobacter pomorum* (*A. pomorum*), *Enterococcus faecalis* (*E. Faecalis*) or *Micrococcus luteus* (*M. Luteus*) failed to trigger activation of this lipogenic regulator (Fig. 1a). These results were confirmed using the transcription of the SREBP target gene, *Acetyl-CoA synthase* (*ACS*) as a readout ^22,26^. *ACS* mRNA levels were increased in *E.cc*-fed flies compared to sucrose-fed flies and this increase was even stronger with *E. coli* (Fig. 1c).

**Figure 1.**
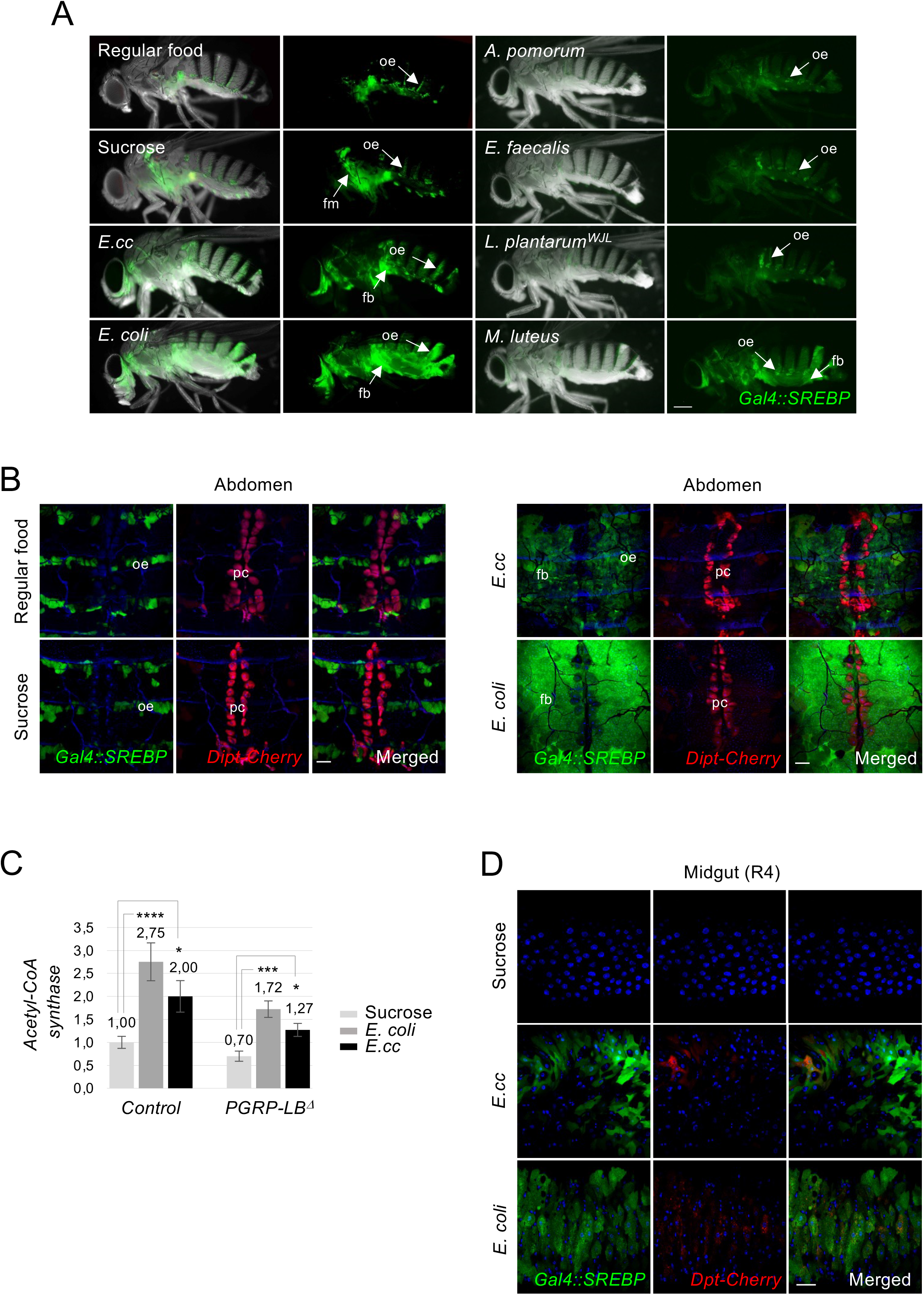
Ingestion of *E. coli* or *E. cc* bacteria activates SREBP both in adipocytes and enterocytes. (A) Pictures of adult flies fed 8 days with either regular food or sucrose or on a mixture of sucrose + bacteria such as *E. cc* or *E. coli* or *A. pomorum* or *E. faecalis* or *L. plantarum^WJL^* or *M. luteus*, and showing *Gal4::SREBP* activation (green). Flies fed on regular food and or sucrose show activation of *Gal4::SREBP* in oenocytes and in flight muscle + oenocytes, respectively. *E. coli* feeding, and to a less extend *E. cc* ingestion, promote activation of *Gal4::SREBP* in fat bodies while ingestion of either *A. pomorum* or *E. faecalis* or *L. plantarum^WJL^* or *M. luteus* do not. Flies of the following genotypes were used: *w^1118^/w^1118^; Gal4::SREBP, UAS-2XEGFP/+*. oe: oenocytes, fm: flight muscles. (B) Confocal images of the dorsal part of adult abdominal carcasses viewed from inside, showing *Dipt-Cherry* expression (red) and *Gal4::SREBP* activation (green) from flies fed on regular food, sucrose or on a mixture of sucrose + bacteria (*E. cc* or *E. coli*). Both *E. coli* and *E. cc* feeding activate *Gal4::SREBP* in fat body cells, while no expression of *Dipt-Cherry* is visible, as expected. Constitutive expression of *Dipt-Cherry* in pericardial cells and activation of *Gal4::SREBP* in oenocytes are indicated. Flies of the following genotypes were used: *w^1118^/w^1118^; Gal4::SREBP, UAS-2XEGFP/+; Dipt-CherryC1/Dipt-CherryC1*). pc: pericardial cells, oe: oenocytes. (C) Histograms showing the expression of *ACS* measured by Q-RT-PCR and performed with mRNA extracted from adult abdominal carcasses of control or *PGRP-LB^Δ^* adults fed 4 days with either sucrose, on with a mixture of sucrose + bacteria (*E. coli* or *E. cc*). The mRNA level in non-infected control flies was set to 1 and values obtained with indicated genotypes were expressed as a fold of this value. Histograms correspond to the mean value ± SD of three experiments. *p<0.05, ***p< 0.001, ****p<0.0001; Kruskal-Wallis test. Flies of the following genotypes were used: *w^1118^/w^1118^;; Dipt-Cherry^C1^/Dipt-Cherry^C1^* (*Control*) and *w^1118^/w^1118^;; PGRP-LB^Δ^, Dipt-Cherry^C1^/PGRP-LB^Δ^, Dipt-Cherry^C1^* (*PGRP-LB^Δ^*). (D) Confocal images of the R4 domain of adult midguts from flies fed with either sucrose or with a mixture of sucrose + bacteria (*E. cc* or *E. coli*), showing *Dipt-Cherry* expression (red) and *Gal4::SREBP* activation (green). Both *E. coli* and *E. cc* feeding activates *Gal4::SREBP* in enterocytes, while a faint expression of *Dipt-Cherry* is induced. Flies of the following genotypes were used: *w^1118^/w^1118^; Gal4::SREBP, UAS-2XEGFP/+; Dipt-Cherry^C1^/Dipt-Cherry^C1^*). Scale bar is 0,25 mm (A), 100 μm (B) and 50 μm (D).

### *E. coli* triggers SREBP activation in enterocytes and ovaries

We then asked whether *E.cc* and *E. coli* would activate lipogenesis in other tissues and organs known to be lipogenic. We first monitored the enterocytes which are in close proximity to the gut microbiota and represent another major source of lipids for the organism. Both *E. coli* and *E.cc-*fed females showed *Gal4::SREBP* activation in midgut enterocytes (Fig. 1d). Since *Drosophila* females have to adapt their metabolic machinery to meet the biosynthetic demands of egg production ^23^, we asked whether these bacteria could also impact lipid metabolism in ovaries. While ovaries of sucrose-fed females were atrophic, those of females raised on an *E. coli*-contaminated solution resemble those of females fed on regular food (Fig. 2a and Supplemental Fig. 3). Consistently, *SREBP* activation was observed in mid-stage follicle of *E. coli* fed females, but not in ovaries from flies fed on sucrose only (Fig. 2b). As expected, if this response was a consequence of the extra energy demand required for offspring production, both gut and fat body of virgin females displayed a much weaker SREBP activation after *E. coli* feeding (Fig. 2c, d). Consistently, males fed with *E. coli* displayed no sign of SREBP activation in enterocyte and only a constitutive no-bacteria dependent activation of SREBP in adipocytes (Fig. 2c, d). Taken as a whole, these data show that gut-associated *E. coli* and *E.cc* can active SREBP-dependent lipogenesis locally in enterocytes and remotely in fat body and ovaries from mated females.

**Figure 2.**
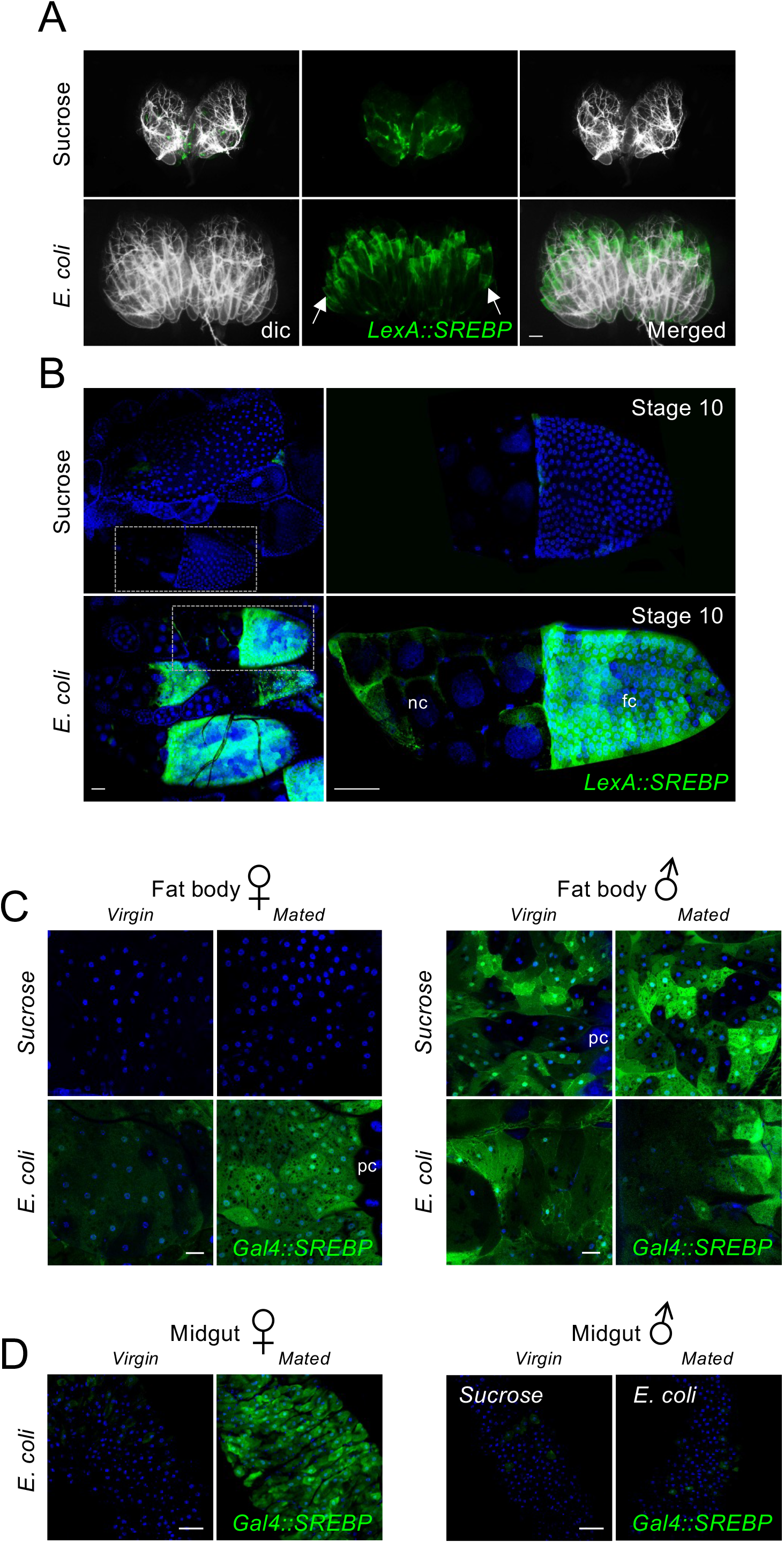
*E. coli* ingestion triggers SREBP activation in ovaries and in fat body of mated females. (A) Pictures of adult ovaries from females fed 2 days with sucrose or with a mixture of sucrose + *E*. *coli*, showing *LexA::SREBP* activation (green). Atrophy of ovaries is obvious in female fed with sucrose, not with *E. coli*. Stage 10 follicles display activation of *LexA::SREBP* (arrows in A), in females fed with *E. coli*. (B) Confocal images of ovarioles dissected from females fed 2 days with sucrose or with *E*. *coli*, showing *LexA::SREBP* activation (green). Females fed with *E. coli*. display activation of *LexA::SREBP* both in nurse cells and in follicle cells of stage 10 egg chambers. nc: nurse cells and fc: follicle cells. (C) Confocal images of fat body from virgin or mated females (or males) fed 2 days on sucrose or on a mixture of sucrose + *E. coli* showing *Gal4::SREBP* activation (green). Activation of *Gal4::SREBP* in adipocytes is strong in mated female and faint in virgin females. Male fat bodies however show no induction but constitutive activation of *Gal4::SREBP* in a *“salt and pepper”* pattern. (D) Confocal images of the R4 domain of adult midguts from virgin or mated females flies fed 2 days with a mixture of sucrose + *E. coli*, or from males flies fed 2 days with sucrose or with a mixture of sucrose + *E. coli*, showing *Gal4::SREBP* activation (green). *E. coli* feeding does not promote activation of *Gal4::SREBP* in enterocytes from virgin females or mated males. Flies of the following genotypes were used: *w^1118^*/*w^1118^, LexA::SREBP*, *13XLexAop2-6XGFP/+* (A-B), *w^1118^*/*w^1118^; Gal4::SREBP, UAS-2XEGFP/+* (females in C-D), *w^1118^*/*Y; Gal4::SREBP, UAS-2XEGFP/+* (males in C-D). pc: pericardial cells, oe: oenocytes. Scale bar is 200 μm (A), 50 μm (B and D) and 20 μm (C and E).

### Bacterial ingestion promotes insulin signaling pathway

Amino-acids sensing by the fat body remotely controls the release of insulin like peptides that increase insulin signaling in peripheral tissue and trigger lipid storage by adipocytes ^27^. To test whether bacteria-mediated SREBP activation corresponds to a modification of the nutritional status the fly, we first monitored insulin signaling in fat body cells using *tGPH* membrane recruitment as a readout. Flies fed with *E.cc* or *E. coli,* or raised on yeast extract as a medium containing AA source, showed *tGPH* membrane recruitment in adipocytes (Fig. 3a). Such an effect was not observed when flies where fed on sucrose only. These results, suggesting that gut *E. coli* and *E.cc* activate insulin signaling in adipocytes, were confirmed using q-RT-PCR on adult’s abdominal carcasses. mRNA levels of the negatively regulated insulin pathway target gene *4EBP/Thor*, were decreased following *E. coli* and *E.cc* feeding (Fig. 3b). We then wondered whether insulin signaling was required for SREBP activation by gut-associated microbiota using *chico^1^*, a loss-of-function allele of the Insulin Receptor Substrate Chico/IRS. We found that, in addition to their expected small size, *chico^1^* mutant females did not show any sign of *LexA::SREBP* activation when fed with *E. coli* (Fig. 3c). The typical signal of *LexA::SREBP* activation in enterocytes and in adipocytes of *E. coli* fed flies was absent in *chico^1^* mutants (Fig. 3d, e). Only a weak, bacteria-independent, SREPB activation was observed in fat bodies and midguts from *chico^1^* animals (Fig. 3d, e). Consistently, we found that mutants for Foxo, a catabolic transcription factor negatively regulated by the insulin pathway, displayed a consistent activation of SREBP in adipocytes, when raised on regular food (Fig. 3f and Supplemental Fig. 2c). In addition, when using a dose of *E. coli* (10 times less concentrated), that is normally not sufficient to activate *Gal4::SREBP* in wild-type flies, *foxo* mutants displayed a clear activation of *Gal4::SREBP* (Fig. 3f).

**Figure 3.**
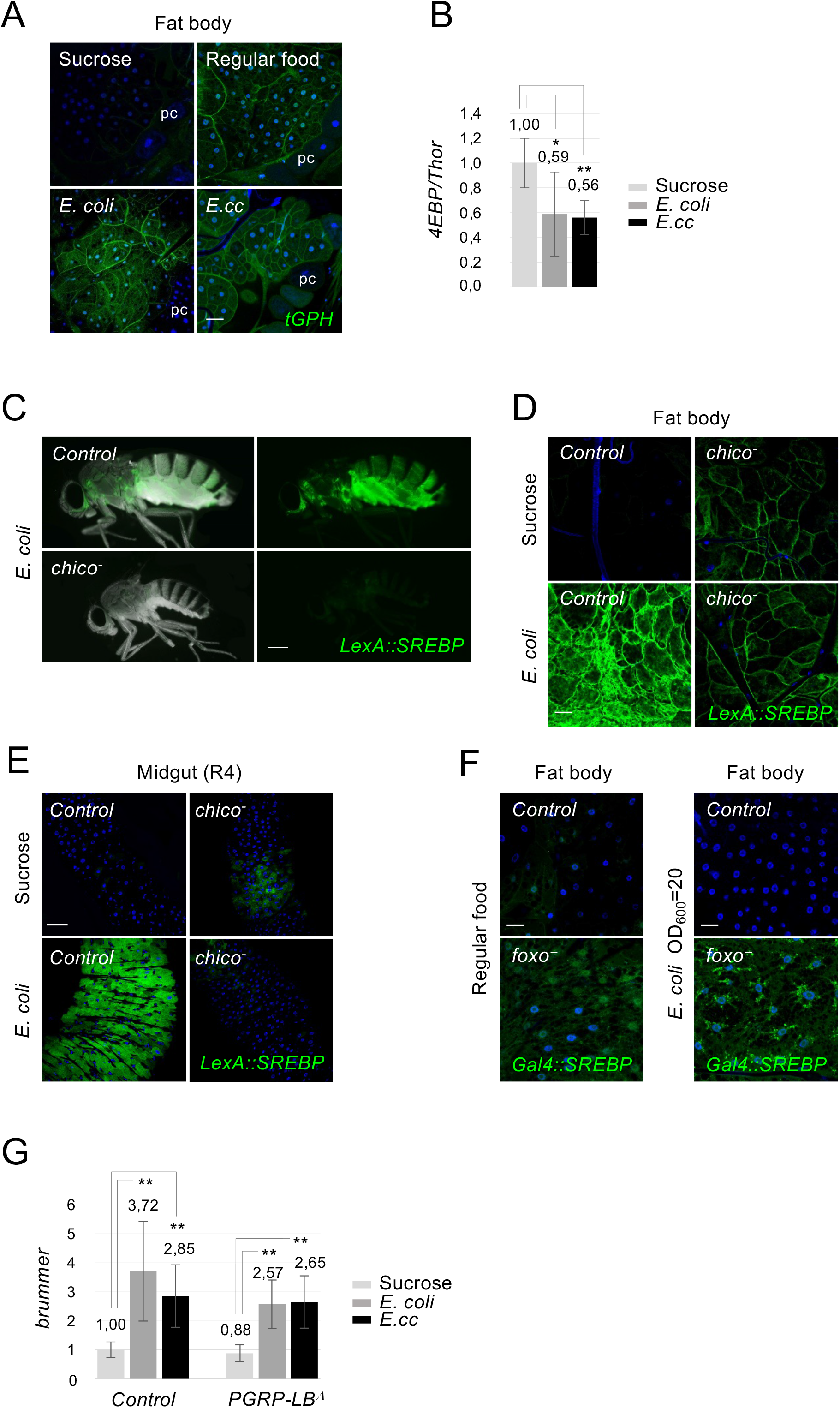
*E. coli* and *E. cc* ingestion promotes systemic insulin signaling pathway and insulin signaling is required for SREBP activation by *E. coli.* (A) Confocal images of fat body from flies fed 4 days with either sucrose, regular food or a mixture of sucrose + *E. coli* or *E. cc*. The expression of the *tGPH* marker is shown in green. An intense recruitment of *tGPH* at the cell surface of adipocytes is observed in flies fed on regular food or orally infected with either *E. coli* or *E. cc*, when compared to a sucrose diet. Flies of the following genotypes were used: *w^1118^*/*w^1118^; tGPH/tGPH*. pc: pericardial cells. (B) Histograms showing the activation of the insulin signaling pathways measured by Q-RT-PCR of *4EBP* mRNAs extracted from adult abdominal carcasses of control adults fed 4 days with either sucrose, on with a mixture of sucrose + bacteria (*E. coli* or *E. cc*). mRNA level in non-infected control flies was set to 1 and values obtained with indicated genotypes were expressed as a fold of this value. Histograms correspond to the mean value ± SD of three experiments. *p<0.05, **p< 0.01; Kruskal-Wallis test. Flies of the following genotypes were used: *w^1118^/w^1118^;; Dipt-Cherry^C1^/Dipt-Cherry^C1^*). (C) Pictures of control or *chico-* mutant flies fed 2 days with *E. coli* showing *LexA::SREBP* activation. Inhibition of insulin signaling pathway abolishes activation of *LexA::SREBP*. (D-E) Confocal images of fat body (D) or midgut R4 domain (E) from control or *chico-* mutant flies fed 2 days with sucrose or with *E. coli*, showing *LexA::SREBP* activation. *E. coli* feeding does not promote activation of *LexA::SREBP* in adipocytes or enterocytes in absence of functional insulin signaling pathway. Flies of the following genotypes were used: *w^1118^*/*w^1118^; LexA::SREBP/Df(2L)ED729; 13XLexAop2-mcd8-GFP/+* (*Control* in C, D and E) and *w^1118^*/*w^1118^; LexA::SREBP, chico1/Df(2L)ED729; 13XLexAop2-mcd8-GFP/+* (*chico-* in C, D and E). (F) Confocal images of fat body from control or *foxo^-^* mutant flies fed 5 days on regular food (left panels) or fed 2 days with a low dose (10x dilution) of *E. coli* (right panels), showing *Gal4::SREPB* activation. Foxo is a negative regulator of SREBP activation in adipocytes. Flies of the following genotypes were used: *w^1118^*/*w^1118^; Gal4::SREBP, UAS-2XEGFP/+; foxo^Δ94^/+* (*Control* in F) and *w^1118^*/*w^1118^; Gal4::SREBP, UAS-2XEGFP/+; foxo^Δ94^/foxo^25^* (*foxo^-^* in F). (G) Histograms showing the expression of *brummer* measured by Q-RT-PCR and performed with mRNA extracted from adult abdominal carcasses of control adults fed 4 days with either sucrose, on with a mixture of sucrose + bacteria (*E. coli* or *E. cc*). The mRNA level in non-infected control flies was set to 1 and values obtained with indicated genotypes were expressed as a fold of this value. Histograms correspond to the mean value ± SD of three experiments. **p< 0.01; Kruskal-Wallis test. Flies of the following genotypes were used: *w^1118^/w^1118^;; Dipt-Cherry^C1^/Dipt-Cherry^C1^* (*Control*) and *w^1118^/w^1118^;; PGRP-LB^Δ^, Dipt-Cherry^C1^/PGRP-LB^Δ^, Dipt-Cherry^C1^* (*PGRP-LB^Δ^*). Scale bar is 20 μm (A), 0,25 mm (C), 20 μm (D and F) and 50 μm (E).

Since gut bacteria-derived PGN can activate NF-κB/Relish in fat body cells and that Relish has been shown to restrain the transcription of the ATGL/Brummer lipase in the same cells^10^, we asked whether the bacteria-dependent lipid accumulation could be due to Brummer repression. In contrast to what was expected if it was the case, *Brummer* transcript levels were increased in the presence of bacteria and followed a regulation that resembles that of *ACS* (Fig. 3g and Fig. 1c). This demonstrated that lipid accumulation in fat body of bacteria fed flies is not secondary to a reduced rate of their degradation by lipase but requires a functional insulin signaling pathway.

### IMD signaling inhibits SREBP cell-autonomously in adipocytes

Although both *E. coli* and *E. cc* are able to activate fat body lipogenesis, we noticed that the effects were stronger with *E. coli* than with *E. cc*. Interestingly, previous works has shown that gut *E. cc* is a stronger inducer of fat body NF-κB signaling than *E. coli*, a difference attributed to the ability of *E. cc* to release PGN in larger amounts than *E. coli*. We hence hypothesized that bacteria-dependent gut-born PGN could buffer *SREBP* activation in fat body cells. To test this hypothesis, we analyzed the effects of gut bacteria colonization in a mutant for PGRP-LB, an enzyme that cleaves PGN into non-immunogenic muropeptides. In such mutants, an excess of gut-born PGN reaches the different immune competent tissues leading to a higher NF-κB pathway activation. Fat body SREBP activation, as well as *ACS* mRNA levels, were weaker in *PGRP-LB* mutants than in wild type controls infected by *E. coli* or *E.cc* (Fig. 4a and Fig.1c). This weaker SREBP activation was paralleled by a stronger NF-κB activation monitored with the *Dipt-mCherry* transgene (Fig. 4a) or by qRT-PCR (Fig. 4b). This was, however, not the case in enterocytes (Fig. 4c). These results were confirmed using isoform specific alleles of the PGN cleaving enzyme *PGRP-LB*. Inactivation of the extracellular isoform (PGRP-LB^PC^ named here PGRP-LB^sec^), which is expected to trigger an increase of circulating PGN levels, lead to an NF-κB signaling upregulation and a reduction of SREBP activation in fat body (Fig. 4e). Such effects were not observed in flies carrying mutations in the cytosolic isoform (PGRP-LB^PD^ named here PGRP-LB^intra^) which do not affect the levels of circulating PGN. Moreover, the lack of *Gal4::SREBP* activation observed in *E. coli* or *E. cc*-fed *PGRP-LB^Δ^* mutant flies, was reverted by the simultaneous inactivation of the IMD pathway core component *Dredd^F64^* (Fig. 5a, b) demonstrating that an excessive IMD signaling can antagonize bacteria-dependent SREBP activation.

**Figure 4.**
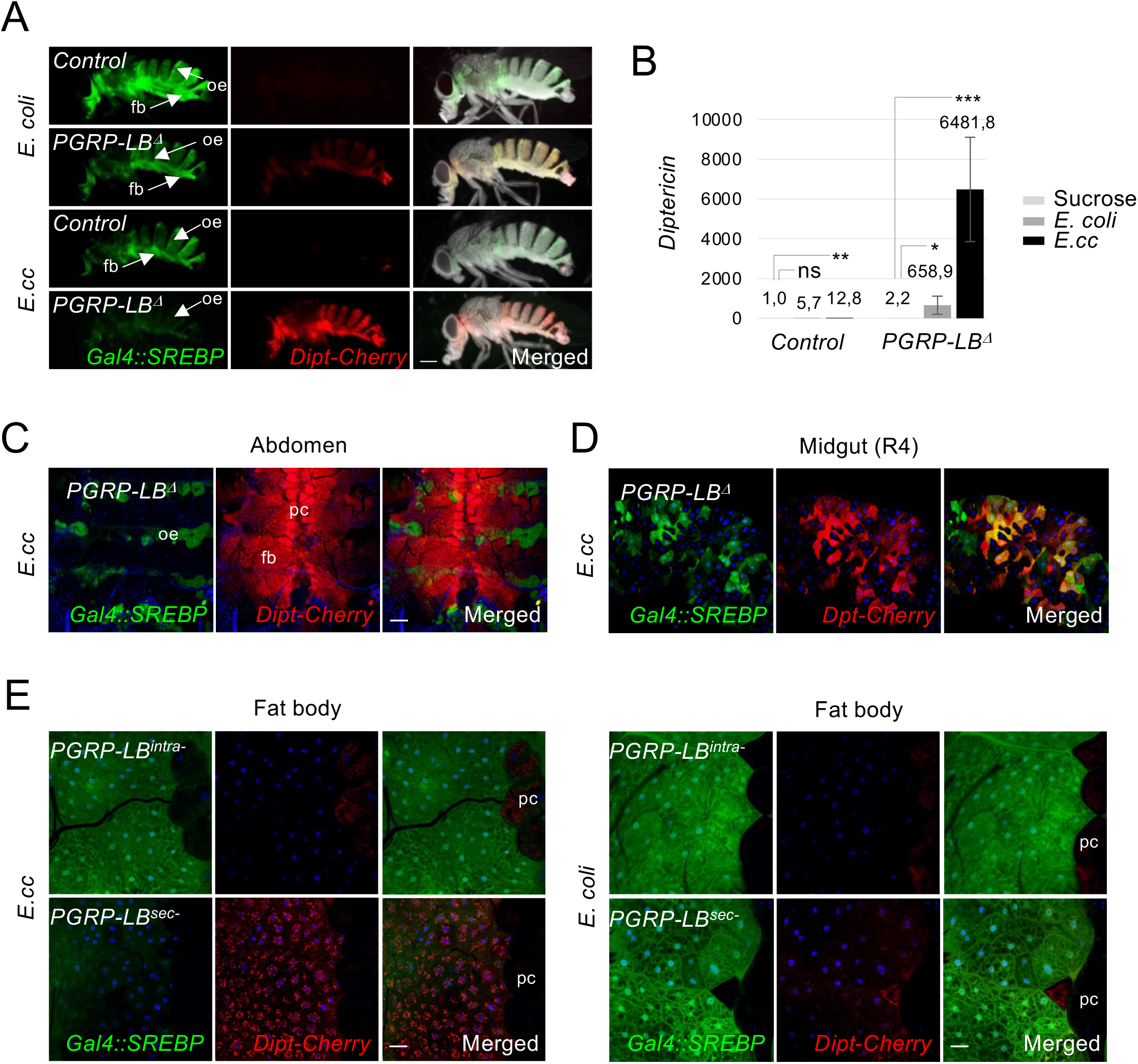
Bacteria-dependent gut-born PGN antagonizes *SREBP* activation in adipocytes. (A) Pictures of adult flies, control or *PGRP-LB^Δ^* mutants, fed 8 days with a mixture of sucrose + *E. coli* or *E. cc*, and showing *Dipt-Cherry* expression (red) and *Gal4::SREBP* activation (green). *E. coli* feeding promotes activation of *Gal4::SREBP* in fat bodies from control and *PGRP-LB^Δ^* mutant. Ingestion of *E. cc*, however, triggers activation of *Gal4::SREBP* in fat body from control flies, but not from *PGRP-LB^Δ^* mutant’s flies. As expected, activation of *Dipt-Cherry* is observed in fat body from *PGRP-LB^Δ^* mutants fed with bacteria. The constitutive activation of *Gal4::SREBP* in oenocytes is indicated (oe arrows). Flies of the following genotypes were used: *w^1118^/w^1118^; Gal4::SREBP, UAS-2XEGFP/+; Dipt-Cherry^C1^/Dipt-CherryC1*(*Control*) and *w^1118^/w^1118^ Gal4::SREBP, UAS-2XEGFP/+; Dipt-Cherry^C1^, PGRP-LB^Δ^/Dipt-Cherry^C1^, PGRP-LB^Δ^* (*PGRP-LB^Δ^*). (B) Histograms showing the expression of *Diptericin* measured by Q-RT-PCR and performed with mRNA extracted from adult abdominal carcasses of control adults fed 4 days with either sucrose, on with a mixture of sucrose + bacteria (*E. coli* or *E. cc*). The mRNA level in non-infected control flies was set to 1 and values obtained with indicated genotypes were expressed as a fold of this value. Histograms correspond to the mean value ± SD of three experiments. *p<0.05, **p< 0.01, ***p<0.001; Kruskal-Wallis test Flies of the following genotypes were used: *w^1118^/w^1118^;; Dipt-Cherry^C1^/Dipt-Cherry^C1^* (*Control*) and *w^1118^/w^1118^;; PGRP-LB^Δ^, Dipt-Cherry^C1^/PGRP-LB^Δ^, Dipt-Cherry^C1^* (*PGRP-LB^Δ^*). (C-D) Confocal images of the dorsal part of adult abdominal carcasses viewed from inside (C) or of the midgut R4 domain (D) from *PGRP-LB^Δ^* mutant flies fed with *E. cc*, showing *Dipt-Cherry* expression (red) and *Gal4::SREBP* activation (green). Adipocytes from *PGRP-LB^Δ^* mutant flies display high level of *Dipt-Cherry* expression but no activation of *Gal4::SREBP*. Constitutive expression of *Dipt-Cherry* in pericardial cells and activation *Gal4::SREBP* in oenocytes are indicated (C). Enterocytes of *PGRP-LB^Δ^* mutant flies display activation of both reporters (D). Flies of the following genotypes were used: *w^1118^/w^1118^ Gal4::SREBP, UAS-2XEGFP/+; Dipt-Cherry^C1^, PGRP-LB^Δ^/Dipt-Cherry^C1^, PGRP-LB^Δ^* (*PGRP-LB^Δ^*). (E) Confocal images of adult fat body from CRISPR mutant flies *PGRP-LB^intra-^* or *PGRP-LB^sec-^*, fed 72h with *E. cc* (A) or *E. coli* (B) and showing *Dipt-Cherry* expression (red) and *Gal4::SREBP* activation (green). (A) *E. cc* feeding induces activation of *Dipt-Cherry* in adipocytes from *PGRP-LB^sec-^* animals, but not from *PGRP-LB^intra-^* ones. *Gal4::SREBP* activation is faint in the CRISPR-specific mutant allele *PGRP-LB^sec-^*and strong in the *PGRP-LB^intra-^*one. (B) *E. coli* feeding induced a comparable activation of *Gal4::SREBP* in both *PGRP-LB^sec-^*or *PGRP-LB^intra-^* adipocyte’s mutant flies, but no activation of *Dipt-cherry*. Flies of the following genotypes were used: *w^1118^/w^1118^;; PGRP-LB^PD*Z10e^, Dipt-Cherry^C1^/PGRP-LB^PD*Z10e^, Dipt-Cherry^C1^* (*PGRP-LB^intra-^*) and *w^1118^/w^1118^;; PGRP-LB^PC*10A^, Dipt-Cherry^C1^/PGRP-LB^PC*10A^, Dipt-Cherry^C1^* (*PGRP-LB^sec-^*). pc: pericardial cells. Scale bar is 0,25 mm (A), 100 μm (C), 50 μm (D) and 20 μm (E).

**Figure 5.**
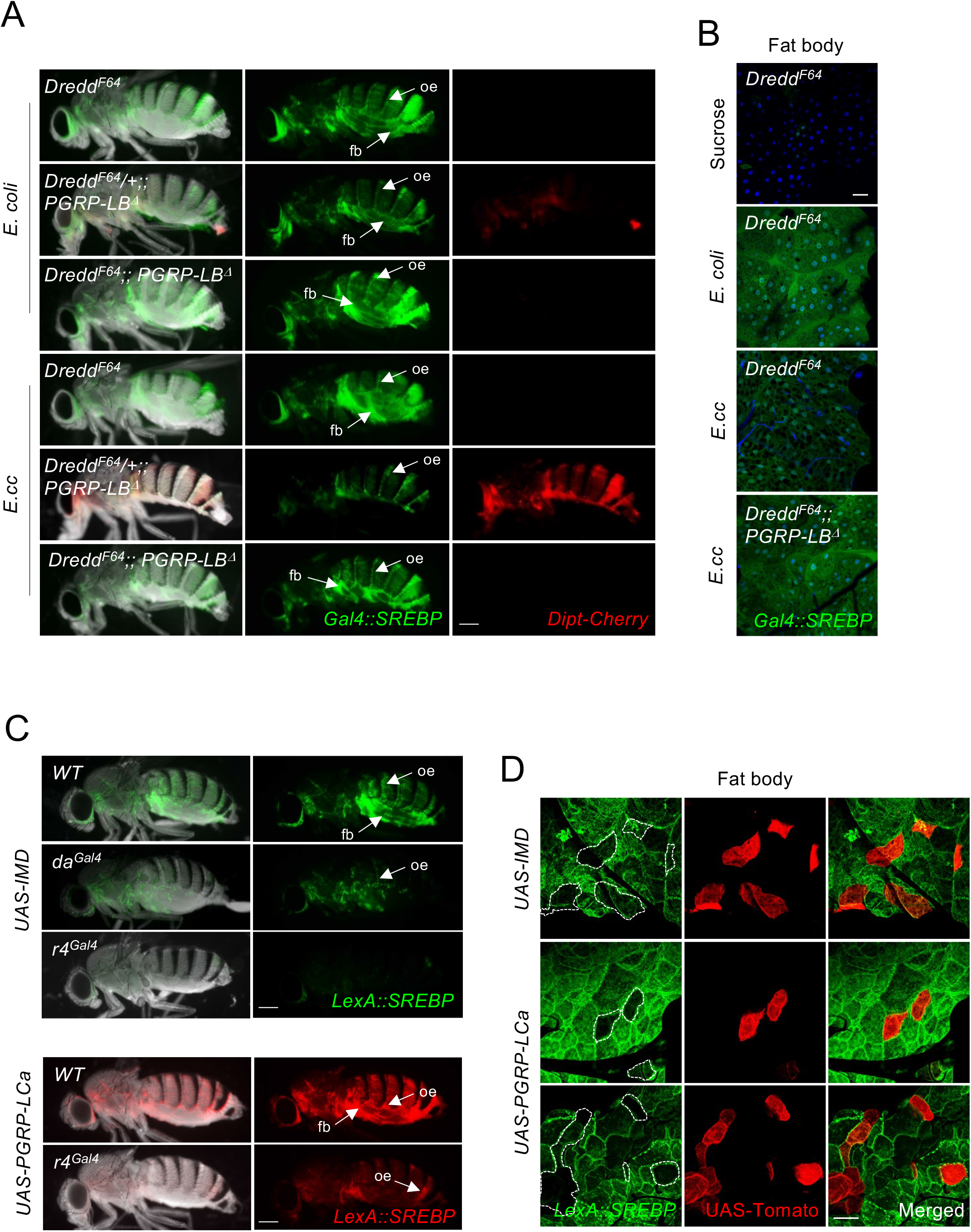
The NF-κB signaling pathway inhibits SREBP activation cell-autonomously in adipocytes. (A) Pictures of adult flies fed 7 days with either *E. coli* (A) or *E. cc* (B) showing *Dipt-Cherry* expression (red) and *Gal4::SREBP* activation (green). Flies mutant for the loss-of-function allele *Dredd^F64^* activate *Gal4::SREBP* in fat body, when fed with either *E. coli* or *E. cc*. Two copies of the *Dredd^F64^* allele suppress the faint and the lack of activation of *Gal4::SREBP* observed in *PGRP-LB^Δ^* mutant flies fed with *E. coli* and *E. cc*, respectively (A). (B) Confocal images of fat body from flies fed 2 days with sucrose or with a mixture of sucrose + the indicated bacteria. Activation of *Gal4::SREBP* (shown in green) absent in sucrose fed flies, is similarly detectable in *Dredd^F64^* mutant flies fed with either *E. coli* or *E. cc*. Double mutant *Dredd^F64^;; PGRP-LB^Δ^* flies activate *Gal4::SREBP* in adipocytes when fed 2 days with *E. cc*. Flies of the following genotypes were used: *w^1118^, Dredd^F64^*/ *w^1118^, Dredd^F64^; Gal4::SREBP, UAS-2XEGFP/+; Dipt-Cherry^C1^/Dipt-Cherry^C1^* (for *Dredd^F64^*) or *w^1118^, Dredd^F64^/ w^1118^; Gal4::SREBP, UAS-2XEGFP/+; Dipt-Cherry^C1^, PGRP-LB^Δ^/Dipt-Cherry^C1^, PGRP-LB^Δ^* (for *Dredd^F64^*/+*;; PGRP-LB^Δ^*) or *w^1118^, Dredd^F64^*/ *w^1118^, Dredd^F64^; Gal4::SREBP, UAS-2XEGFP/+; Dipt-Cherry^C1^, PGRP-LB^Δ^/Dipt-Cherry^C1^, PGRP-LB^Δ^* (for *Dredd^F64^;; PGRP-LB^Δ^*). (C) Pictures of adults flies fed 3 days with *E. coli* showing *LexA::SREBP* activation (green in top panels and red in bottom panels). The activation of *LexA::SREBP* is inhibited in fat bodies from flies over expressing IMD or PGRP-LCa in fat body cells with *r4^Gal4^*, or from flies over expressing IMD ubiquitously with *da^Gal4^*. Flies of the following genotypes were used: *w^1118^, w^1118^*/*w^1118^, LexA::SREBP*, *13XLexAop2-6XGFP/+; r4Gal4/+* (*Control* in top panels) or *w^1118^*/*w^1118^, LexA::SREBP*, *13XLexAop2-6XGFP/+; daGal4 or r4Gal4/UAS-IMD* (*da^Gal4^* and *r4^Gal4^* in top panels), or *w^1118^*/*w^1118^, LexA::SREBP*, *LexAop-mCherry.mito/+; r4Gal4/+* (*Control* in bottom panels) or *w^1118^*/*w^1118^, LexA::SREBP*, *LexAop-mCherry.mito/+; r4Gal4/UAS-PGRP-LCa* (*r4^Gal4^* in bottom panels). (B) Confocal images of fat body showing clones of adipocytes overexpressing IMD or PGRP-LCa (red) and activation of *LexA::SREBP* (green), from flies fed 2 days with *E. coli*. Fat body clones over expressing either IMD or PGRP-LCa inhibits *LexA::SREBP* activation is a strictly autonomous manner. Flies of the following genotypes were used: *w^1118^*, *CoinFLP^Gal4^*/*w^1118^*; *LexA::SREBP, LexAop-CD8-GFP-2A-CD8-GFP, UAS-CD4::Tomato/hs-FLP.G5, Tub^Gal80ts^; UAS-IMD or UAS-PGRP-LCa/+.* Scale bar is 0,25 mm (A and C) and 20 μm (B and D).

To identify the tissue(s) in which NF-κB activation is required for SREBP regulation, we monitored the effects of IMD pathway components over expression, which is sufficient to activate downstream signaling in the absence of bacteria. *PGRP-LCa* or *IMD* overexpression, either ubiquitously (*da^Gal4^*) or specifically in the fat body (*r4^Gal4^*), prevented the activation of *LexA::SREBP* in *E. coli* fed flies (Fig. 5c). Moreover, clonal over expression of IMD or *PGRP-LCa* cell-autonomously prevented *LexA::SREBP* activation in fat body of females fed with *E. coli* (Fig. 5d). These data demonstrate that the microbiota-dependent SREBP activation in fat body cells is cell-autonomously repressed by a PGN-dependent IMD/NF-κB pathway activation.

### Inhibition of IMD signaling in adipocytes improves survival of *E. cc* infected *PGRP-LB^Δ^*

To test the physiological relevance of this antagonism, we monitored the survival curves and lipid droplet accumulation in various genetic combinations chronically infected with *E. cc*. As expected, whereas wild-type flies fed with *E. cc* showed lipid droplet accumulation, this was not the case for *PGRP-LB* mutants (Fig. 6a). In addition, *PGRP-LB* mutants died much earlier than their wild-type siblings upon *E.cc* chronic infection (Fig. 6b). As shown in for SREBP activation (Fig. 5a), the reduced lifespan and lipid droplet non-accumulation in adipocytes observed in *E.cc*-infected *PGRP-LB^Δ^* mutants were restored by the simultaneous inactivation of IMD pathway component *Dredd* (Fig. 6a, b). Since IMD/NF-κB signaling specifically inhibits SREBP activation and lipogenesis in adipocytes, we tested whether IMD signaling buffering in adipocytes could ameliorate the survival of *PGRP-LB^Δ^* mutant flies chronically infected with *E.cc*. To do so, we took advantage of a *UAS-dFadd^IR^* transgene whose targeted expression can block IMD signaling over activation typically observed in guts and fat bodies of *E. cc*-infected *PGRP-LB^Δ^* mutant (Fig. 6c). We then tested the effects of a tissue-specific IMD silencing on the lifespan of chronically *E. cc*-infected *PGRP-LB^Δ^* mutants. We found that blocking IMD signaling in enterocytes with *Mex^Gal4^* or in muscles using *Mef2^Gal4^* did not ameliorate the lifespan of *E. cc* fed *PGRP-LB^Δ^* mutants. In contrast, ubiquitous and fat body specific expression of *UAS-dFadd^IR^* significantly improved *PGRP-LB^Δ^* mutant resistance to chronic *E. cc* infection (Fig. 6d). These results suggest that by buffering IMD pathway activation in the fat body, the PGRP-LB amidase allows this tissue to generate lipids and hence to better resist to chronic infection.

**Figure 6.**
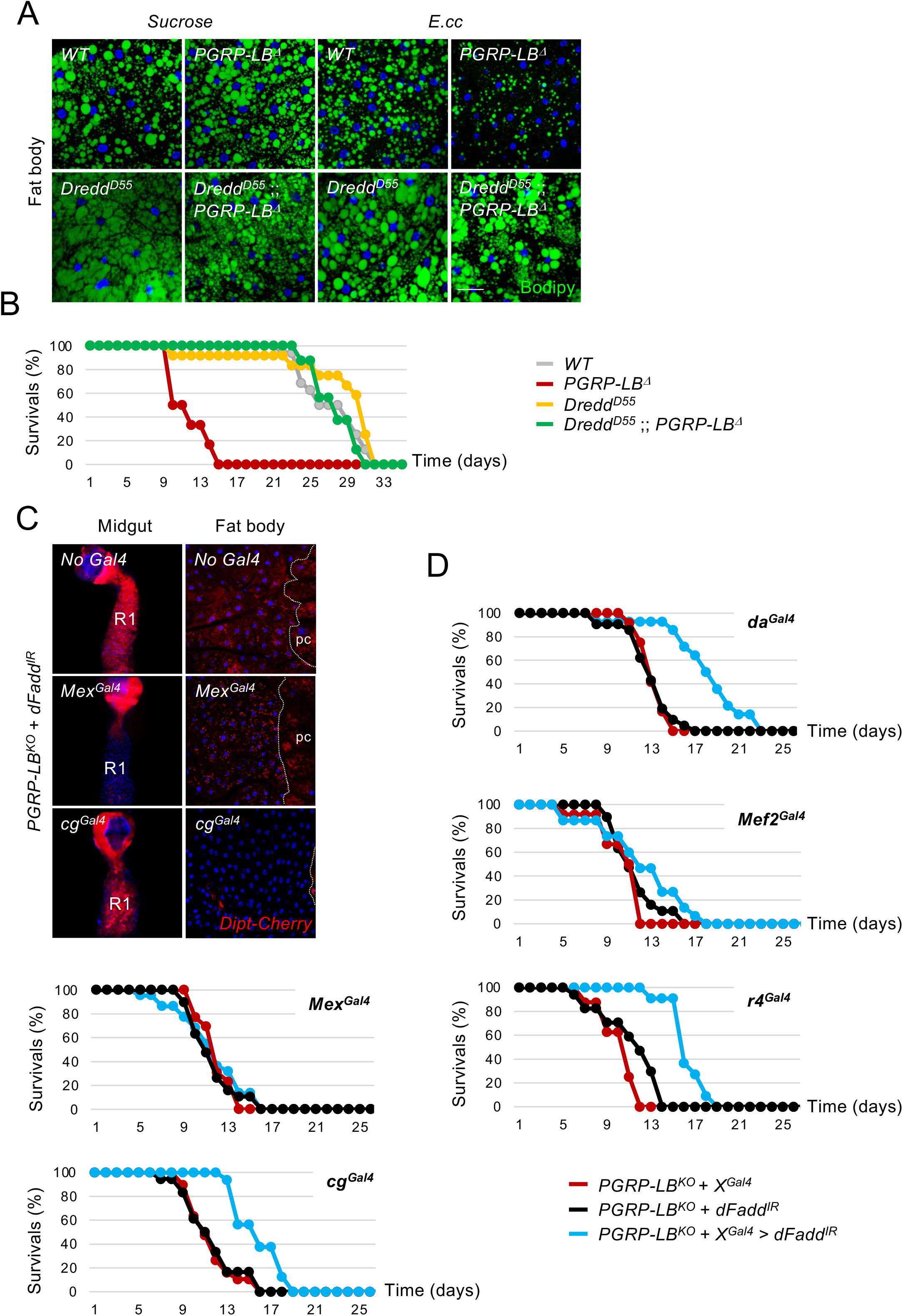
Inhibition of the NF-κB signaling pathway in adipocytes ameliorates survival of *PGRP-LB^Δ^* mutant flies infected with *E. cc*. (A) Confocal images of fat body from flies fed 8 days with sucrose or with a mixture of sucrose + *E. cc*, and showing lipids droplets labelled with Bodipy (green). The reduction of lipid storage characteristic of *E. cc* infected *PGRP-LB^Δ^* flies is suppressed in presence of two copies of the loss of function allele *Dredd^D55^*. (B) Survival analysis of flies orally infected with *E. cc*. The *Dredd^D55^* allele suppresses the deleterious effect of *PGRP-LB^Δ^* mutation on fly’s survival. For A and B, flies of the following genotypes were used: *w^1118^/w^1118^* (*Control*), *w^1118^/w^1118^;; PGRP-LB^Δ^/PGRP-LB^Δ^* (*PGRP-LB^Δ^*), *w^1118^, Dredd^D55^/w^1118^, Dredd^D55^* (*Dredd^D55^*) and *w^1118^, Dredd^D55^/w^1118^, Dredd^D55^;; PGRP-LB^Δ^/PGRP-LB^Δ^* (*Dredd^D55^;; PGRP-LB^Δ^*). (C) Confocal images of the R1 domain of adult midgut (left panels) or of adult fat body (right panels) from *PGRP-LB^Δ^* mutant fed 24h with *E. cc*, showing *Dipt-Cherry* (red) expression. Inhibition of the IMD signaling pathway via expression of *UAS-dFadd^IR^* is effective in enterocytes or in adipocytes using *Mex^Gal4^* or *cg^Gal4^*, respectively. (D) Survival analysis of *PGRP-LB^Δ^* mutant flies orally infected with *E. cc*. The expression of *UAS-dFadd^IR^* in adipocytes, using either *cg^Gal4^* or *r4^Gal4^* decreases the deleterious effect of *PGRP-LB^Δ^* mutation on fly’s survival. Flies of the following genotypes were used: *w^1118^*/*w^1118^; +/+* or *MexGal4/+* or *cg^Gal4^/+; PGRP-LB^Δ^, Dipt-Cherry^C1^*/*PGRP-LB^Δ^, UAS-dFadd^IR^* (C-D) and *w^1118^*/*w^1118^;; PGRP-LB^Δ^, da^Gal4^* or *PGRP-LB^Δ^, Mef2^Gal4^* or *PGRP-LB^Δ^, r4^Gal4^*/*PGRP-LB^Δ^, UAS-dFadd^IR^* (D). Scale bar is 100 μm (midguts in C) and 20 μm (fat bodies in A and C).

**Figure 7.**
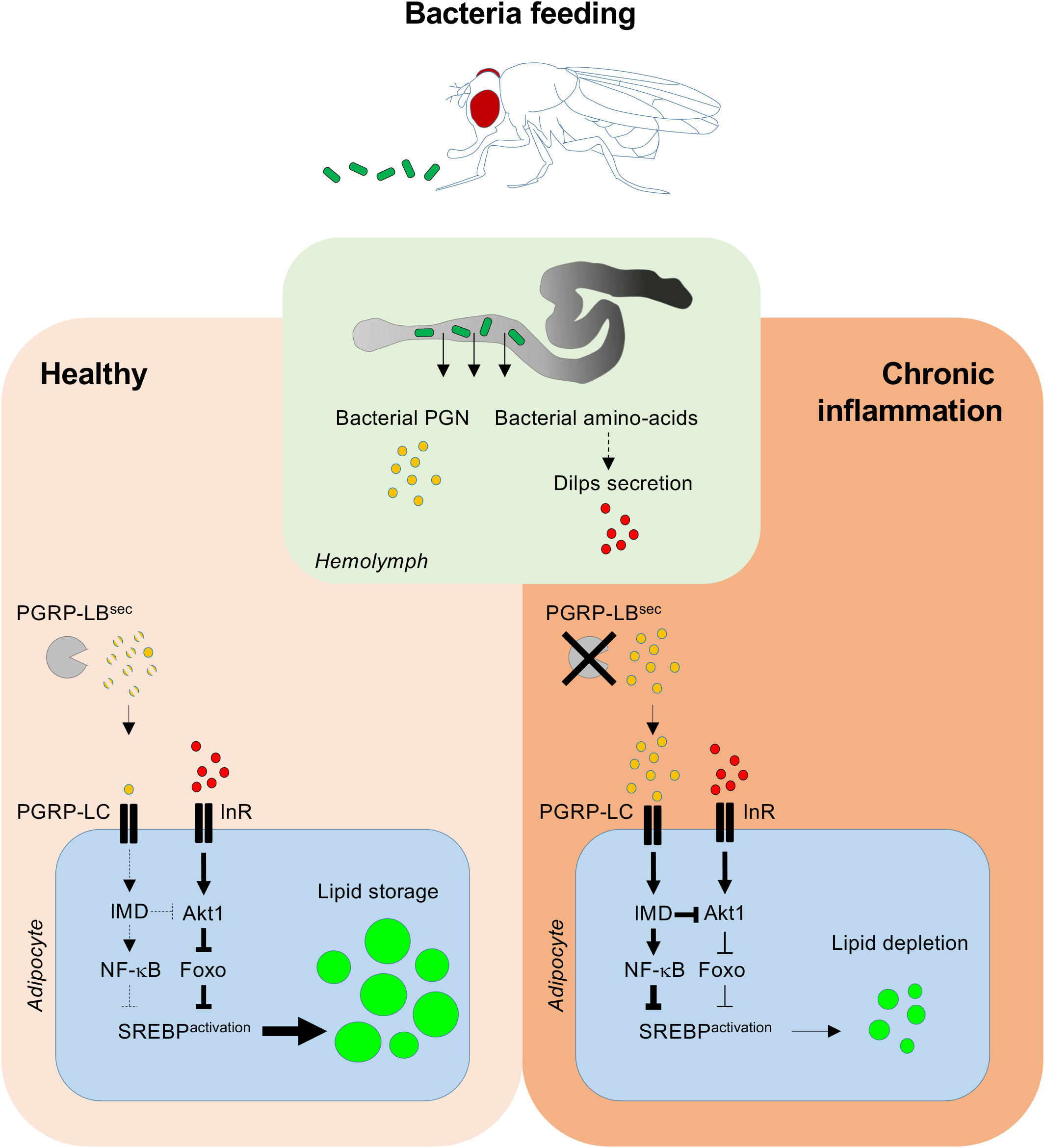
Model for the role of bacterial PGN and bacterial amino-acids in lipid storage formation. In healthy flies, feeding bacteria under protein scarcity promotes bacterial amino-acids transfer from the gut into the hemolymph. This indirectly triggers Dilps secretion by neurosecretory cells of the central brain (not shown), leading to the systemic activation of the insulin/Tor signaling pathways and the activation of SREBP in adipocytes in order to sustain lipogenesis. Simultaneously, gut-born PGN diffuses into the hemolymph where it is degraded by the secreted amidase PGRP-LB^sec^. Upon chronic inflammation due to the lack of PGRP-LB^sec^ and accumulation of PGN, the constitutive activation of the IMD signaling pathway promotes lipid depletion via inhibition of SREBP activation in adipocytes.

## Discussion

Although it is well accepted that gut microbiota affects host energy balance and contributes to the onset and maintenance of metabolic disease(s), the underlying mechanisms are of multiple origins and, in most cases, remains unclear ^28–30^. We showed here that gut-associated bacteria can influence host lipid metabolism by activating SREBP in adipocytes. Axenic flies fed with sucrose displayed phenotypes of undernourished animal, such as ovarian atrophy and reduced systemic insulin signaling. At contrary, *E. coli* fed animal had fully developed ovaries and displayed local (gut) and systemic (fat body) activation of the insulin signaling, genetically upstream of SREBP activation in these lipogenic organs. Which bacteria associated metabolites or constituents mediate these effects? Dietary amino-acids that are required for the release of insulin-like peptides and insulin receptor activation are obvious candidates ^27,31^. Composed of more than 50% of proteins, bacteria could potentially serve as a source of dietary amino-acids. Consistently, while live yeast (*Issatchenkia orientalis*) promotes amino-acids flux from the diet to the host, inactivated ones improve undernourished flie’s lifespan when added to the diet ^32^. Likewise, fly microbiota contributes to protein processing upstream of the nutrient-sensing Tor and insulin signaling pathways to promote systemic growth ^33,34^. In addition, we recently reported that starved adult flies, but not well-fed siblings, favored a bacteria-contaminated sucrose solution over an axenic one, in a two-choice feeding assay ^35^. These data suggest that in case of protein scarcity, flies can use microbes as amino-acids source leading to the activation of the insulin/Tor signaling pathways. This is consistent with previous studies suggesting that insects can use microbes as a food source ^32,36,37^. In this context, SREBP activation in fat tissue could be used as an indicator of an efficient nutrient flux from the microbiota to the host.

Among the microbial strains tested, only *E. coli* and *E. cc* were able to activate the lipogenic program in fly’s adipocytes. Ingestion of *L. plantarum^WJL^*, *A. pomorum*, *E. faecalis*, *M. luteus or Saccharomyces cerevisiae* (data not shown) did not activate SREBP in adipocytes. This lack of effect is consistent with previous results showing that neither *S. cerevisiae* nor *L. plantarum* are able to efficiently rescue adult’s lifespan of undernourished flies ^32^. Further investigation will be necessary to uncover the strains specific compounds and mechanism(s) responsible for this species-specific regulation of SREBP and, in a broader aspect, to stimulate lipid metabolism.

The insulin/Tor signaling pathway controls cell growth and metabolism by boosting protein biosynthesis and lipogenesis, in part via Akt-dependent activation of SREBP ^38^. In mammals, insulin positively regulates SREBP1c and promotes lipogenesis in hepatocytes ^39^. In this line, we found that activation of SREBP in both enterocytes and adipocytes, upon bacteria feeding, was abolished in flies lacking a functional insulin signaling pathway. Surprisingly however, axenic flies fed on a regular diet did not activate SREBP even though the systemic insulin pathway was on. This suggests that activation of insulin signaling pathway *per se* is not sufficient to promote SREBP-dependent lipogenesis in adipocytes. Since SREBP processing is negatively regulated by sterols in mammals and by palmitate or its derivative in *Drosophila* ^40,41^, lipids contained in the fly food ingredients (yeast extract and cornmeal flour) could prevent SREBP processing. In support of this hypothesis we found that flies feeding on a regular food supplemented with *E. coli* displayed a faint and heterogenous activation of *Gal4::SREBP* in adipocytes, compared to flies fed on a Sucrose + *E. coli* diet (Supplementary Fig. 4a).

Other signaling molecules such as sex hormones might contribute to SREBP activation by gut bacteria. Local production of the steroid hormone ecdysone by oocytes follicle cells stimulates SREBP processing, leading to *LpR2* transcription in nurse cells ^23^. By binding to lipoprotein particles circulating in the hemolymph, LpR2 regulates oocyte lipid uptake. Feeding flies with *E. coli* was sufficient to promotes SREBP activation in oocytes, and possibly, to induce ecdysone synthesis by follicle cells. In response to mating, another steroid hormone, the Juvenile hormone induces SREBP activity in enterocytes ^24^. We found that the activation of SREBP in adipocytes and enterocytes by gut bacteria was specifically observed in mated female, and is reliant on the sex peptide receptor SPR (Supplementary Fig. 4b). Several reports connect the sex peptide (SP) to the corpus allatum, an endocrine gland responsible for JH production ^42,43^, suggesting that the systemic effects of mating via SP could be carried out through this pathway. In response to gut microbiota, a female specific endocrine network involving JH, ecdysone and insulin, could stimulate SREBP activation both in gut and fat body cells in order to anticipate an increased lipid requirement.

In *Drosophila*, immunity and metabolism are linked structurally via the fat body, an organ homologous to the mammalian liver, adipose tissue and immune system, made of a single cell type: the adipocyte ^44^. In mammals, immune cells are embedded into the adipose tissue, allowing direct influence of one cell type on the other ^7^. Our data indicate that chronic activation of the IMD/NF-κB pathway prevents gut bacteria-dependent SREBP processing and thus lipid metabolism. By restricting the diffusion of PGN to the fly hemolymph, the PGRP-LB^sec^ enzyme allows microbiota-dependent lipogenesis in remote adipocytes and promote fly survival. In the absence of such brake, lipid storages of orally infected flies are rapidly depleted and life span is reduced. Since *E. cc*-fed *PGRP-LB^Δ^* mutant display ovarian atrophy associated with a reduction of vitellogenic stages ^45^, it is possible that of sex-hormones mis-regulation is contributing to SREBP processing inhibition in this organ.

We found that Foxo is a negative regulator of SREBP processing in adipocytes and that IMD/NF-κB signaling pathway inhibits SREBP processing, without affecting Insulin/PI3K signaling (Supplementary Fig. 5b). Thus, we propose that the NF-κB transcription factor Relish and Foxo acts in parallel, or together, to negatively regulates SREBP processing. Interestingly, both transcription factors have common immune and metabolic target genes in *Drosophila* fat body ^6,10,46,47^. One possibility would be that Relish and Foxo negatively regulate the transcription of genes that are essentials for SREBP processing, such as the escort factor SCAP (<u>S</u>REBP <u>C</u>leavage <u>A</u>ctivating <u>P</u>rotein), and/or the proteases S1P (<u>S</u>ite-<u>1</u> <u>P</u>rotease) and S2P (<u>S</u>ite-<u>2</u> <u>P</u>rotease) ^26,48^.

Inhibition of lipid metabolism triggered by bacterial infection have been reported in the past, although in different contexts. When bacteria such as *Mycobacterium marinum* are injected into *Drosophila* body cavity, the transcription factor Mef2, which activates transcription of metabolic genes in non-infected individuals, switches its activity to enhance transcription of immune genes ^8^. As a result, anabolic transcripts are reduced and energy stores, such as lipids, are lost. Toll and the IMD signaling pathways are acting genetically upstream of Mef2 in this process. Lee and colleagues found that *E. cc* infection triggers lipid catabolism in enterocytes which, via a TRAF3-AMPK/WTS-ATG1 pathway, contributes to the activation of DUOX, a member of the NADPH oxidase family acting as the first line of host defense in *Drosophila* gut ^49^. Finally, the bacteria produced short chain fatty acid acetate acts as a microbial metabolic signal that activates signaling through the IMD pathway in enteroendocrine cells. This, in turn, increases transcription of the endocrine peptide tachykinin, which is essential for timely larval development and optimal lipid metabolism and insulin signaling ^50^.

Our work sheds light on how gut microbiota influences lipid metabolism and contributes to the development of an immune-metabolic disorder, through the action of the highly conserved transcription factors SREBP, NF-κB and Foxo and the universal bacteria cell-wall component PGN. Furthermore, it shows how by buffering microbiota-born circulating PGN levels, the PGRP-LB amidase, allows the appropriate balance between metabolic and immune responses.

## Acknowledgments

This work was supported by (ANR-11-LABX-0054) (Investissements d’Avenir–Labex INFORM), ANR PEPTIMET (ANR-18-CE15-0018-02), Equipe Fondation pour la Recherche Médicale (EQU201903007783) and Institut Universitaire de France to J.R

---

**Supplementary Figure 1.**
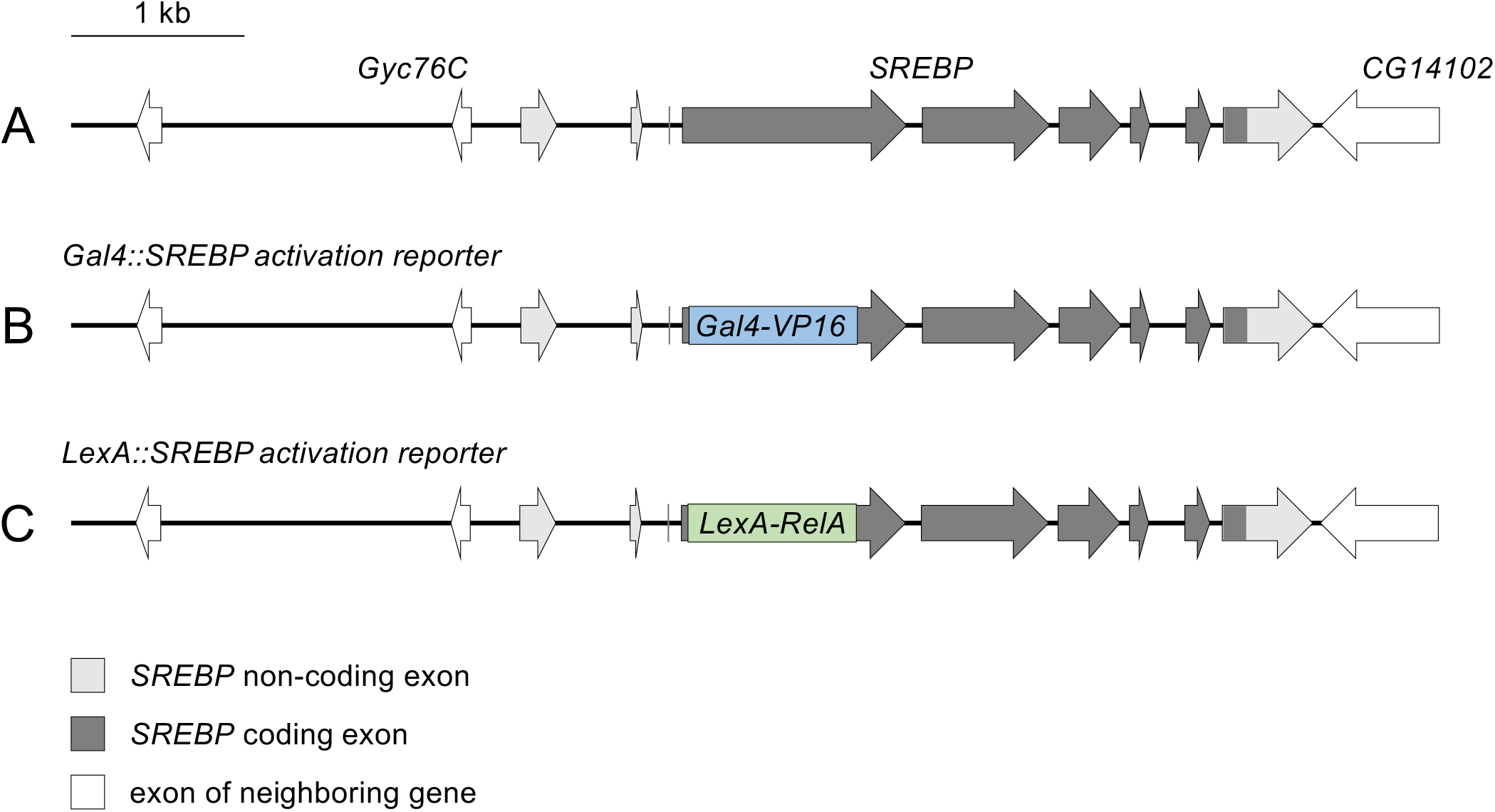
*Gal4::SREBP* and *LexA::SREBP* transgenes used in this study. 1, Schematic drawing of the *SREBP* genomic locus. 2, In the *Gal4::SREBP* transgene, the transcription factor domain-encoding sequence was replaced by a Gal4::VP16-encoding sequence to report SREBP activation. 3, In the *LexA::SREBP* transgene the transcription factor domain-encoding sequence was replaced by a LexA::RelA-encoding sequence to report SREBP activation.

**Supplementary Figure 2.**
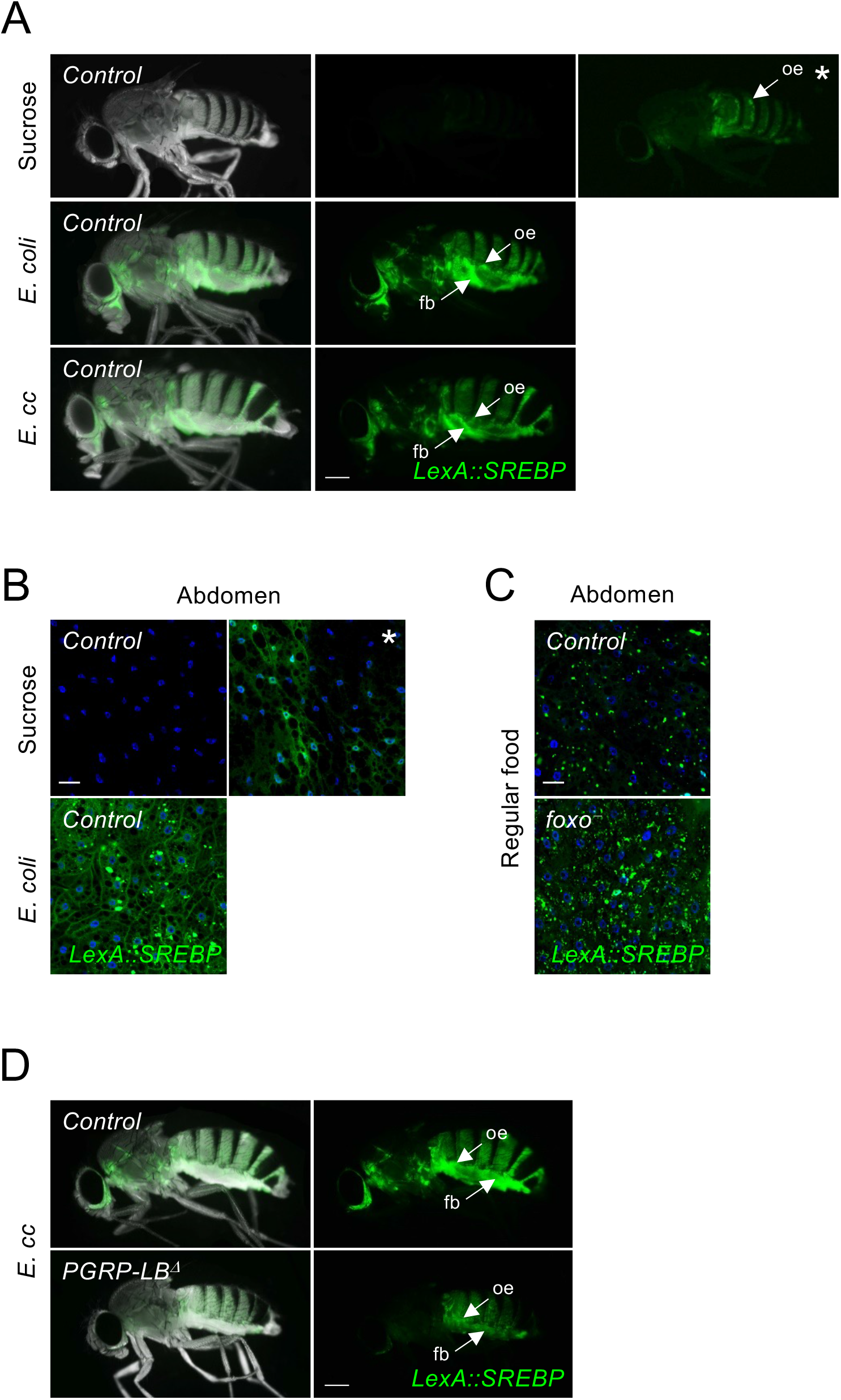
Gut bacteria stimulate LexA::SREBP activation in adult fat body. (A) Pictures of adult flies fed 2 days with sucrose, or a mixture of sucrose + *E. coli* or *E. cc*, showing *LexA::SREBP* activation (green). Flies fed with sucrose show activation of *LexA::SREBP* in oenocytes, noticeable after lowering the signal detection threshold (panel with asterisk). Both *E. coli* and *E. cc* feeding promotes activation of *LexA::SREBP* in fat bodies. (B) Confocal images of fat body from flies fed 2 days with sucrose or with *E. coli* and showing *LexA::SREBP* activation (green). Flies fed on sucrose show feeble activation of *LexA::SREBP* in adipocytes, noticeable after lowering the signal detection threshold (panel with asterisk). *E. coli* feeding, however, promotes strong activation of *LexA::SREBP* in adipocytes. (D) Pictures of adult flies, control or *PGRP-LB^Δ^* mutant, fed 2 days with sucrose + *E. cc*, showing *LexA::SREBP* activation (green). Ingestion of *E. cc* triggers activation of *LexA::SREBP* in fat body from control flies, but not from *PGRP-LB^Δ^* mutant’s flies. Flies of the following genotypes were used: *w^1118^*/*w^1118^, LexA::SREBP*, *13XLexAop2-6XGFP/+* (*Control* in A, B and D), *w^1118^*/*w^1118^, LexA::SREBP*, *13XLexAop2-6XGFP/+; foxo^Δ94^/+* (*Control* in C), *w^1118^*/*w^1118^, LexA::SREBP*, *13XLexAop2-6XGFP/+; foxo^Δ94^/foxo^25^* (*foxo^-^* in C) and *w^1118^*/*w^1118^, LexA::SREBP*, *13XLexAop2-6XGFP/+; PGRP-LB^Δ^/PGRP-LB^Δ^* (*PGRP-LB^Δ^* in D). Scale bar is 0,25 mm (A and D) and 20 μm (B and C).

**Supplementary Figure 3.**
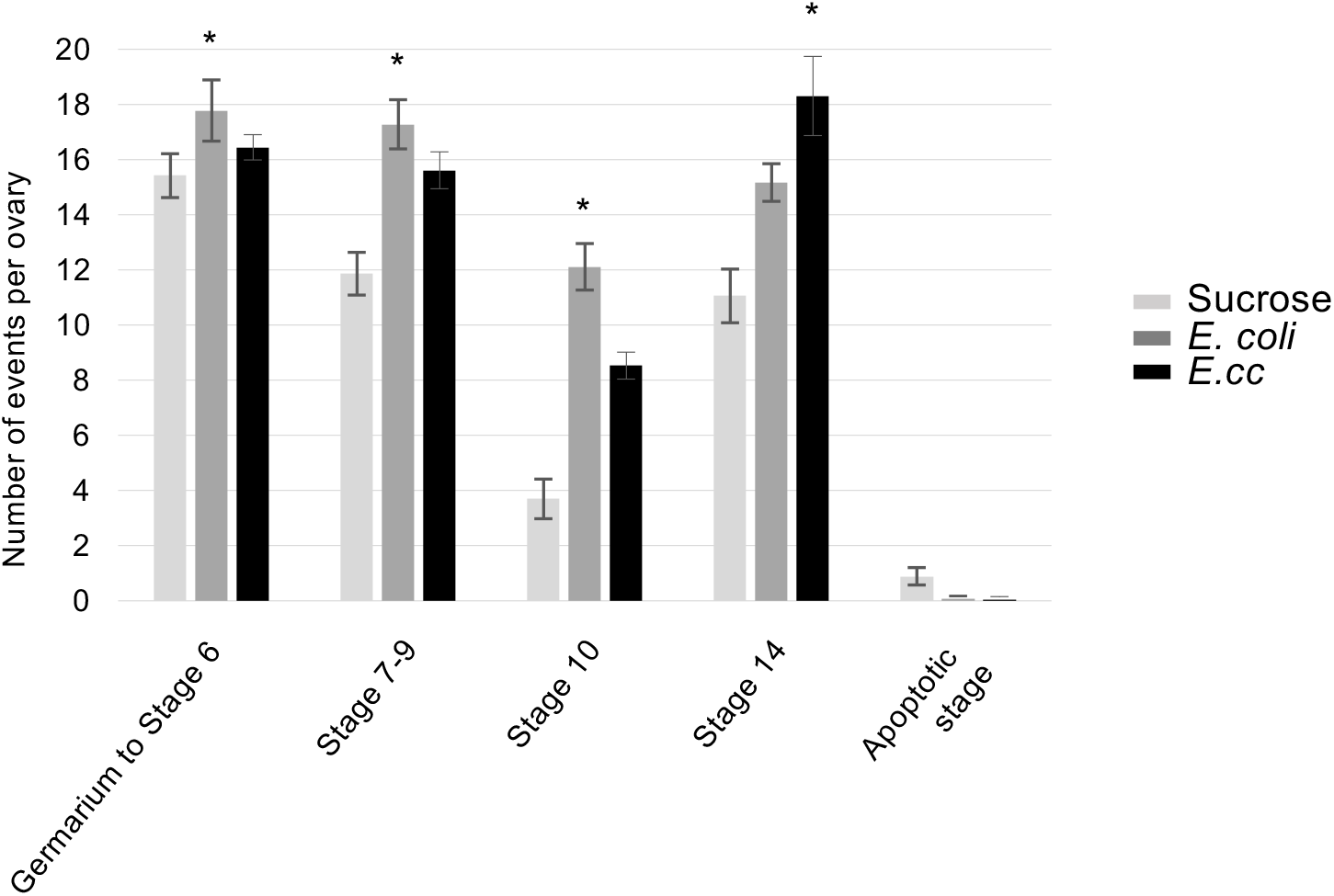
Gut microbiota sustains oogenesis. Quantification of the different stages of oocytes observed in female’s ovary, after feeding 24h on sucrose, or on a mixture of sucrose + *E. coli* or *E. cc*. Apoptotic events were quantified as oocytes with compact and dense nurse cell nuclei, using DAPI staining (not shown). Histograms correspond to the mean value ± SD of three experiments. For each oocyte stage, sucrose values were used as reference for statistical analysis. *p<0.05; Kruskal-Wallis test. Flies of the following genotypes were used: *w^1118^*/*w^1118^; LexA::SREBP*, *13XLexAop2-6XGFP/+*.

**Supplementary Figure 4.**
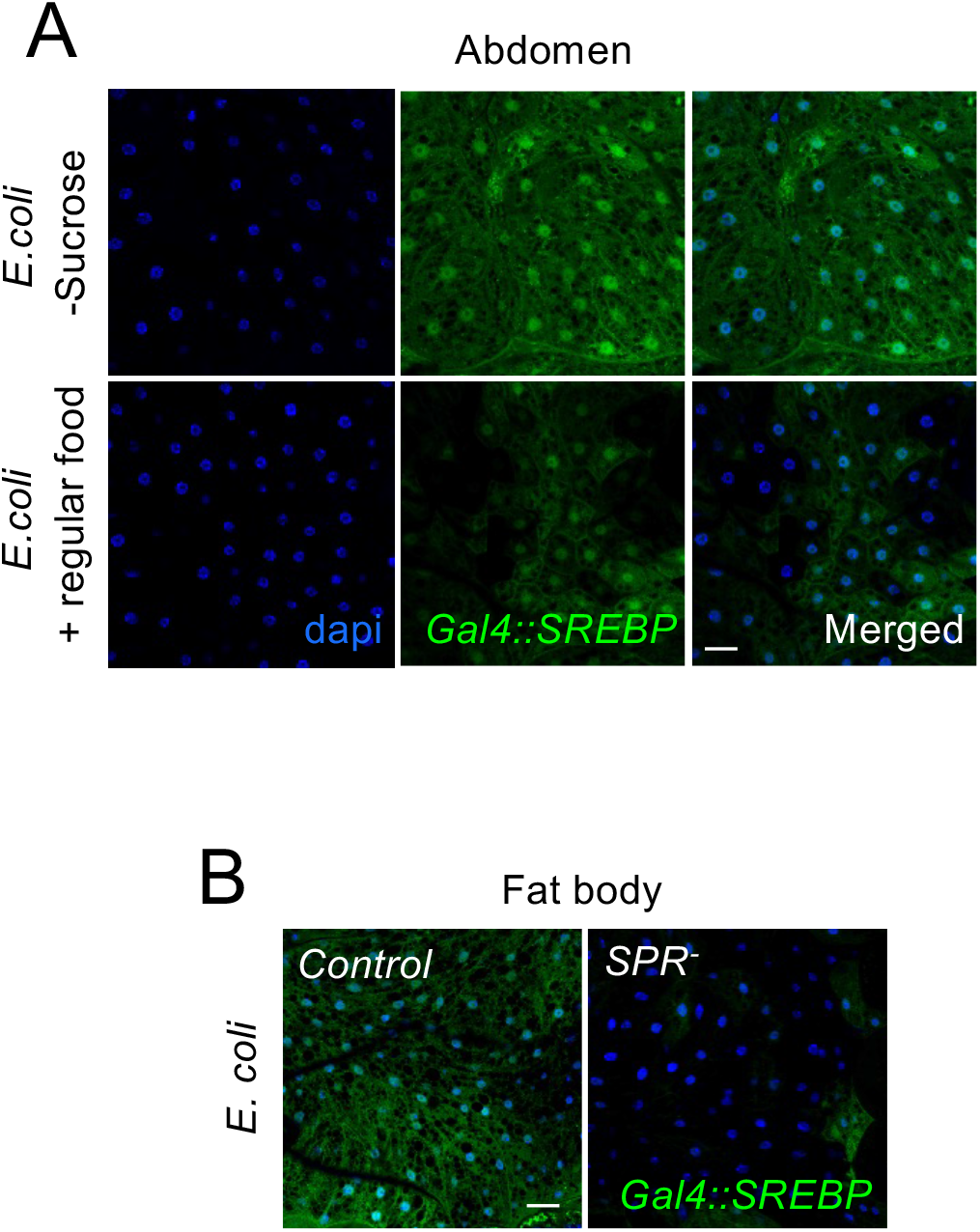
Activation of SREBP by *E. coli* on regular food and Sex Peptide Receptor requirement for SREBP activation in adipocytes. (A) Confocal images of fat body from female flies fed 2 days with *E. coli* without sucrose, or with *E. coli* dropped on regular food, showing *Gal4::SREBP* activation (green). Absence of sucrose does not impact the strong activation of SREBP by *E. coli*, while presence of regular food diminishes it. (B) Confocal images of fat body from control or *SPR^-^* mutant females fed 2 days with *E. coli* showing *Gal4::SREBP* activation (green). The sex peptide receptor SPR is required for activation of *Gal4::SREBP* in adipocytes. Flies of the following genotypes were used: *w^1118^*/*w^1118^; Gal4::SREBP, UAS-2XEGFP/+* (A), and *w^1118^, Df(1)SPR/w^1118^; Gal4::SREBP, UAS-2XEGFP/+* (*Control* in B) and *w^1118^, Df(1)SPR/ w^1118^, Df(1)SPR; Gal4::SREBP, UAS-2XEGFP/+* (*SPR^-^* in B). Scale bar is 20 μm.

**Supplementary Figure 5.**
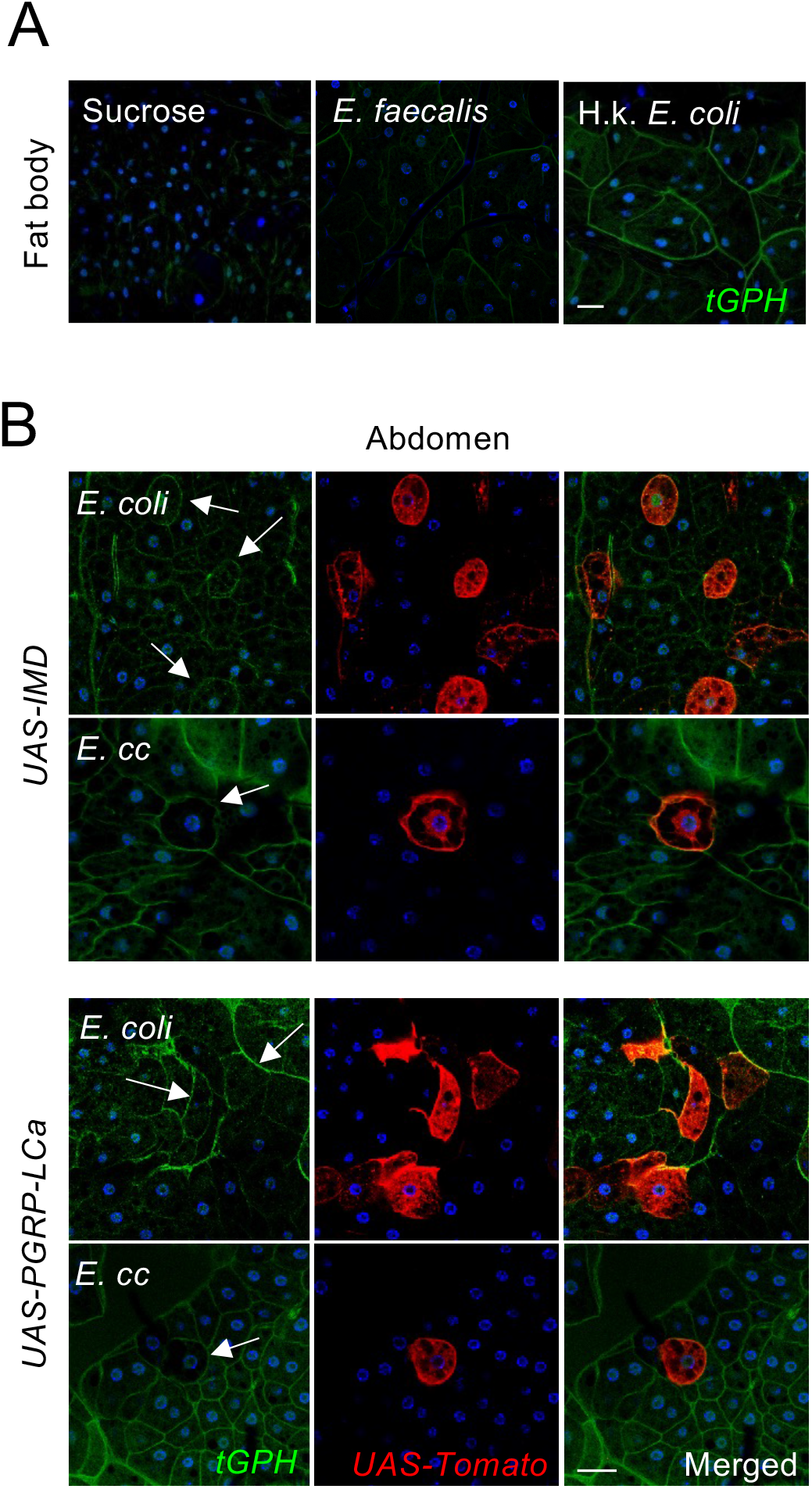
Gut microbiota promote cell surface recruitment of tGPH in adipocytes and this is not affected by over expressing IMD or PGRP-LCa cell autonomously. (A) Confocal images of fat body from adult flies fed 2 days with either sucrose, or a mixture of sucrose + *E. faecalis* or heat killed (H.k) *E. coli*, and showing the *tGPH* marker (green). Ingestion of heat killed *E. coli* promote recruitment of *tGPH* at the cell surface of adipocytes, compared to a sucrose diet. (B) Confocal images of fat body showing clones of adipocytes overexpressing IMD or PGRP-LCa (red) and the *tGPH* marker (green), from flies fed 1 day with *E. cc* or 2 days with *E. coli*. Fat body clones over expressing either IMD or PGRP-LCa do not affect *tGPH* recruitment at the cell surface of adipocytes. Flies of the following genotypes were used: *w^1118^*/*w^1118^; tGPH/tGPH* (A) and *w^1118^*, *CoinFLP^Gal4^*/*w^1118^*; *tGPH, UAS-CD4::Tomato/hs-FLP.G5, Tub^Gal80ts^; UAS-IMD or UAS-PGRP-LCa/+.* Scale bar is 20 μm.

## Material and methods

### *Drosophila* strains and maintenance

The strains used in this work are: *w^1118^* BL#3605, *Gal4::SREBP* from BL#38395, *UAS-2XEGFP* BL#6874, *Diptericin-Cherry^C1^* ^51^, *PGRP-LB*^*Δ* 21^, *Dredd^F64^* (a gift from François Leulier), *da^Gal4^* BL#55851, *r4^Gal4^* and *Mex^Gal4^* (kindly provided by Yixian Zheng), *Mef2^Gal4^* BL#27390, *cg^Gal4^* BL#7011, *UAS-IMD* (kindly provided by François Leulier), *UAS-PGRP-LCa* BL#30917, *CoinFLP^Gal4^* BL#59269, *hs-FLP.G5* BL#58356, *Tub^Gal80ts^* BL#7108, *UAS-CD4::Tomato* (kindly provided by Frank Schnorrer), *LexA::SREBP* (this work, molecular details of the construct under request), *13XLexAop2-6XGFP* BL#52265, *LexAop-CD8-GFP-2A-CD8-GFP* BL#66545, *tGPH* BL#8163, *13XLexAop2-mcd8-GFP* BL#32203, *chico^1^* BL#10738, *Df(2L)ED729* BL#24134, *foxo^Δ94^* BL#42220, *foxo^25^* BL#80944, *Df(1)SPR* BL#7708, *PGRP-LB^PD*Z10e^* and *PGRP-LB^PC*10A^* (Kurz et al., 2017), *UAS-dFadd^IR^* (^52^; kindly provided by Pascal Meier). Flies were grown at 25°C on a yeast/cornmeal medium in 12h/12h light/dark cycle-controlled incubators. For 1liter of food, 8.2g of agar (VWR, cat. #20768.361), 80g of cornmeal flour (Westhove, Farigel maize H1) and 80g of yeast extract (VWR, cat. #24979.413) were cooked for 10 min in boiling water. 5.2 g of Methylparaben sodium salt (MERCK, cat. #106756) and 4 ml of 99% propionic acid (CARLOERBA, cat. #409553) were added when the food had cooled down. For antibiotic (ATB) treatment, the standard medium was supplemented with Ampicillin, Kanamycin, Tetracyclin and, Erythromycin at 50 μg/ml final concentrations.

### *Drosophila* genetics and analysis

To generate *UAS-IMD* and *UAS-PGRP-LCa* overexpressing clones, 5 days old mated females were raised and aged in presence of ATB at 22°C. Adult flies were then transferred into non-ATB media, and placed 24h at 29°C to inactivate Gal^80ts^, before the infection by bacteria for the following 48h at 29°C. No heat shock was required for clone induction. Flies of the following genotype were used: *CoinFLP^Gal4^*/+*; hs-FLP.G5/LexA::SREBP*, *LexAop-CD8-GFP-2A-CD8-GFP; UAS-IMD, Tub^Gal80ts^ /+ or PGRP-LCa, Tub^Gal80ts^ /+* for Figure 5D or *CoinFLP^Gal4^*/+*; hs-FLP.G5/tGPH, UAS-CD4::Tomato; UAS-IMD, Tub^Gal80ts^ /+ or PGRP-LCa, Tub^Gal80ts^ /+* for Supplementary Figure 4B.

### Imaging

Whole fly imaging was performed on adult females totally immersed in 70% EtOH. Images were captured using a ZEISS SteREO Discovery.V12 microscope. For dissected tissues, adult flies were cut apart in cold PBS, fixed for 20 min in 4% paraformaldehyde on ice and rinsed 3 times in PBT (1XPBS + 0.1% Triton X-100). The tissues were mounted in Vectashield (Vector Laboratories) fluorescent mounting medium, with or without DAPI. Images were captured with an LSM 780 ZEISS confocal microscope.

### Bodipy and Nile red Staining

For Bodipy and Nile Red staining, adult tissues were dissected in PBS, fixed for 20 min in 4% paraformaldehyde on ice, rinse 3 times in 1XPBS without detergent and stained with Nile red (Cat. No. 72485, Sigma-Aldrich) or Bodipy™ 493/503 (Cat. No. D3922, ThermoFisher) at respectively 1:10000 and 1:1000, in PBS for 30 min.

### Bacterial strains

The following microorganisms were used: *Erwinia carotovora carotovora 15* strain 2141 (grown at 30°C), *Lactobacillus plantarum* strain WJL (grown at 37°C), *Escherichia coli* strain DH5*α* (grown at 37°C), *Acetobacter pomorum* (grown at 30°C), *Enterococcus faecalis* (grown at 37°C) and *Micrococcus luteus* (grown at 30°C). Microorganisms were cultured overnight in Luria-Bertani (for *E. cc*, *E. coli, E. faecalis* and *M. luteus*) or MRS medium (for *L.plantarum* and *A. pomorum*). Cultures were centrifuged at 4000 g for 15 min at RT and re-suspended in 1XPBS. Cells were serially diluted in 1XPBS and their concentration was determined by optical density (OD) measurement at 600 nm.

### Adult oral infection

We used 4-6 days old female raised at 25°C in presence of ATB in the food. 24h before the infection, female flies were transferred in vials without ATB and then placed in a fly vial with microorganism contaminated food. The food solution was obtained by mixing a pellet of an overnight culture of bacteria or yeast (OD=200) with a solution of 5% sucrose (50/50) and added to a filter disk that completely covered the agar surface of the fly vial. For *E. coli* heat inactivation, a solution of *E. coli* diluted (final OD_600_=100) in 2,5% Sucrose was incubated at 96°C for 20 minutes, then cool down before use.

### Survival Tests with Bacterial Infection

For oral infections, adult flies were transferred every 2 days in a fresh vial in which 150 microliters of a fresh solution of *E. cc* (OD =200)/ 5% sucrose (50/50) has been deposited.

### Quantitative Real-Time PCR

RNA from whole dissected organs (n=12) was extracted with RNeasy Mini Kit. Three hundred ng of total-RNA was then reverse transcribed in 10 μl reaction volume using the Superscript III enzyme (Invitrogen) and random hexamer primers. Quantitative real-time PCR was performed on a CFX96 Real-Time PCR Detection System (BIO-RAD) in 96-well plates using the FastStart Universal SYBR Green Master (Sigma-Aldrich). The amount of mRNA detected was normalized to control rp49 mRNA values. Normalized data was used to quantify the relative levels of a given mRNA according to cycling threshold analysis (ΔCt). All datasets were organized and analyzed in Microsoft Excel 2016.

### Statistical Analysis

The Prism software (GraphPad) was used for statistical analyses. We used the nonparametric Kruskal-Wallis test. P value was indicated as follow: * for P<0,05, ** for P<0,01, *** for P<0,001. ns for not significantly different.

